# Machine Learning Maps Research Needs in COVID-19 Literature

**DOI:** 10.1101/2020.06.11.145425

**Authors:** Anhvinh Doanvo, Xiaolu Qian, Divya Ramjee, Helen Piontkivska, Angel Desai, Maimuna Majumder

**Author notes:** Corresponding Authors: Anhvinh Doanvo, Maimuna Majumder.

## Abstract

**Summary:** Manually assessing the scope of the thousands of publications on the COVID-19 (coronavirus disease 2019) pandemic is an overwhelming task. Shortcuts through metadata analysis (e.g., keywords) assume that studies are properly tagged. However, machine learning approaches can rapidly survey the actual text of coronavirus abstracts to identify research overlap between COVID-19 and other coronavirus diseases, research hotspots, and areas warranting exploration. We propose a fast, scalable, and reusable framework to parse novel disease literature. When applied to the COVID-19 Open Research Dataset (CORD-19), dimensionality reduction suggested that COVID-19 studies to date are primarily clinical-, modeling- or field-based, in contrast to the vast quantity of laboratory-driven research for other (non-COVID-19) coronavirus diseases. Topic modeling also indicated that COVID-19 publications have thus far focused primarily on public health, outbreak reporting, clinical care, and testing for coronaviruses, as opposed to the more limited number focused on basic microbiology, including pathogenesis and transmission.

## Introduction

On March 16, 2020, the White House issued a ‘call to action’ for the application of artificial intelligence (AI) methods to assist with research on coronavirus disease 2019 (COVID-19) (Robbins, 2020), which is caused by the severe acute respiratory syndrome coronavirus 2 (SARS-CoV-2), designating machine learning (ML) as a potentially useful tool for gleaning critical insights from the existing coronavirus literature. Present attempts to examine COVID-19-related publications either mine texts to rapidly create study summaries for researchers (Joshi et al., 2020) or focus on citations, keyword co-occurrences, and other metrics to identify influential literature (Chahrour et al., 2020; Golinelli et al. 2020; Hossain, 2020). Other large-scale efforts concentrate on cataloguing peer-reviewed COVID-19 studies, including “LitCOVID”, a literature hub by the National Center for Biotechnology Information (Chen et al., 2020), and “COVID-19 Data Portal”, a literature search engine from the European Bioinformatics Institute (EMBL-EBI, 2020). Although these efforts facilitate keyword-based searches to rapidly identify studies of interest and, in LitCOVID specifically, classify them into broad categories (e.g., mechanism, diagnosis, etc.), they do not provide an overview of where research efforts are directed and whether these efforts have changed over time. This presents an opportunity to leverage the application of ML methods to survey the ongoing influx of peer-reviewed and pre-printed COVID-19 studies – combined with prior publications on severe acute respiratory syndrome (SARS) coronavirus (SARS-CoV), Middle East respiratory syndrome (MERS) coronavirus (MERS-CoV), and other coronaviruses – and develop unique insights for COVID-19 research needs.

A number of biomedical studies have already applied ML techniques in their work on surveillance, trends, and clinical predictors for the ongoing pandemic (e.g., Alimadadi et al., 2020; Carrillo-Larco & Castillo-Cara, 2020; Ge et al., 2020; Kim et al., 2020; Kumar et al., 2020; Rao and Vazquez, 2020; Yan et al., 2020). Our novel application of ML methods to available coronavirus abstracts, including those about COVID-19, offers insights into the themes of COVID-19 research that overlap with studies about other coronaviruses. We perform ML-aided analysis of research abstracts in the COVID-19 Open Research Dataset (CORD-19) (Wang et al. 2020) to automatically categorize ongoing research endeavors into dynamically-generated categories, enabling us to identify topics that have received limited attention to date. By understanding the knowledge overlap between recently released abstracts on COVID-19 and abstracts related to other coronaviruses, we are able to gain insight into potential areas of SARS-CoV-2 research warranting further exploration. In addition, we propose a reusable framework for parsing an existing knowledge base about other emerging pathogens like the highly pathogenic avian influenza H5N1 (Kilpatrick et al., 2006; Kissler et al., 2019) before they escalate to the level of a major epidemic or pandemic threat.

## Materials and Methods

**Without using any pre-existing knowledge about the abstracts’ topics**, we employed unsupervised ML to determine differences between COVID-19 and non-COVID-19 abstracts in our corpus of documents. A dimensionality reduction approach was used to identify principal patterns of variation in the abstracts’ text, followed by topic modeling to extract high-level topics discussed in the abstracts (James et al., 2013). Our data pipeline is available on GitHub^1^.

### Dataset and Preprocessing

We obtained research abstracts from CORD-19 on May 28, 2020. Generated by the Allen Institute for AI, and in partnership with other research groups, CORD-19 is updated daily with coronavirus-related literature. Peer-reviewed studies from PubMed/PubMed Central, as well as pre-prints from bioRxiv and medRxiv, are retrieved using specific coronavirus-related keywords (“COVID-19” OR “Coronavirus” OR “Corona virus” OR “2019-nCoV”OR “SARS-CoV” OR “MERS-CoV” OR “Severe Acute Respiratory Syndrome” OR “Middle East Respiratory Syndrome”). At time of writing, CORD-19 contained approximately 137,000 articles, including both full-text and metadata for all coronavirus research articles, with ∼40% of the dataset classified as virology-related (Wang et al. 2020). We focused our analysis on the abstracts of articles in CORD-19.

As some of the CORD-19 abstracts were neither relevant to SARS-CoV-2 nor other coronaviruses, we filtered the CORD-19 data to isolate coronavirus-specific abstracts by searching for abstracts that mentioned relevant terms. These abstracts served as our “documents” associated with the document-term matrices (DTMs) in our natural language processing (NLP) pipeline (Supplemental Information 1). We also identified abstracts for only COVID-19-related studies by filtering for COVID-19-related keywords within this subset (Supplemental Information 2).

### Methodology

#### Dimensionality Reduction

Principal components analysis (PCA) is a dimensionality reduction algorithm that summarizes data by determining linear correlations between variables (Hotelling, 1933). PCA identifies individual patterns of variance, or principal components (PCs), in DTMs that differentiate documents from one another, highlighting key trends in the data (Supplemental Information 3). For example, in a simple corpus with two mutually exclusive topics, like machine learning and health infrastructure, the terms “machine” and “learning” would be correlated with one another. PCA would recognize these terms as an important source of variation, providing a way to differentiate documents about either topic (“machine learning” vs. “health infrastructure”) by the frequency of these terms.

When PCA is applied to DTMs, PCs represent patterns differentiating different documents, ordered by their prominence. Each detected pattern reflects both the contextual links between words and their level of importance within the texts. Words with component values of the greatest magnitude on each PC most strongly drive the pattern that each individual PC recognizes. For example, if “machine” and “healthcare” respectively have highly negative and highly positive values on a particular PC, then that PC detects the pattern that when “machine” appears in a text, “healthcare” appears less often. Another PC may detect a different pattern of variance, such as when some documents mention “deep learning” more often than others.

The projection values of the text corpus onto the PCs suggest what concept each document discusses and to what extent, relative to the average document within the corpus. Following the previous example, strongly negative projection values on the first PC, which would capture the data’s most prominent patterns, indicate that the document mentions “machine” more often than the average and thus, is more likely to focus on machine learning. In addition, projection values on the second PC could distinguish between machine learning documents by focus, or lack thereof, on deep learning or other techniques. This approach enables us to delineate between different groups of abstracts by visualizing differences in their projections on the top PCs. So in short, after applying PCA to the DTMs of our abstracts, we identified which PCs successfully separated COVID-19 and non-COVID-19 abstracts. We then used the component values with the largest magnitude on these PCs to interpret them.

#### Topic Modeling

After establishing high-level trends using PCA, we used latent dirichlet allocation (LDA), a topic modeling method, to add nuance to observed differences between COVID-19 and non-COVID-19 literature and examine potential topics of interest. LDA is an unsupervised probabilistic algorithm that extracts hidden topics from large volumes of text (Blei, Ng and Jordan, 2003). Once trained to discover words that separate documents into a predetermined number of topics, LDA can estimate the “mixture” of topics associated with each document. These mixtures suggest the dominant topic for a document that is then used to assign a document to an overarching topic category. For example, LDA may separate documents into two topics, one on “machine learning” and another on “healthcare”, and if a particular document’s mixture is 60% “machine learning” and 40% “healthcare”, it would assign that document to a “machine learning” topic category.

The predetermined number of topics is the most important hyperparameter in an LDA model, as models with sub-optimal number of topics fail to summarize data in an efficient manner (Blei, Ng and Jordan, 2003; Zhao et al., 2015). The number of topics can be determined by (1) identifying a model that has a low perplexity score and high coherence value when applied to an unseen dataset or (2) conducting a principled, manual assessment of the topics that arise. Perplexity is a statistical measure of how imperfectly the topic model fits a dataset, and a low perplexity score is generally considered to provide better results (Zhao et al., 2015). Similarly, topic models with high coherence values are considered to offer meaningful, interpretable topics (Aletras et al., 2013; Newman, Bonilla, and Buntine, 2011). Thus, a model with a low perplexity score and a high coherence value is more desirable when choosing the optimal number of topics. Our initial implementation of LDA showed no optimal value for the number of topics, even as it approached ∼100, potentially reflecting a relatively shallow yet broad pool of COVID-19 publications. We ultimately identified 30 topics via manual review of topics from topic models with different numbers of topics to identify which model satisfied two criteria: (1) topics that were relatively specific, focusing on a single subject matter, and (2) topics that would typically be non-redundant with one another.

## Results

Our initial corpus included 137,326 entries in the CORD-19 dataset (as of May 28, 2020). 107,557 entries had abstracts available, and of those, 35,281 entries (26% of 47,928) had abstracts mentioning search terms related to coronaviruses. Those that did not mention coronavirus search terms in their abstracts contained coronavirus-related terms somewhere else in the text, such as in its citations. Of the latter subset, 18412 publications (∼50% of the subset, or ∼13% of the entire CORD-19 dataset) were COVID-19-related publications.

### PCA Indicates Limited Number of Laboratory Studies on Viral Mechanisms of SARS-CoV-2

While PCA highlighted the abstracts’ most prominent patterns in the first PC, these patterns were not effective at distinguishing between COVID-19 and non-COVID-19 literature. Figure 1a demonstrates no meaningful difference between the two distributions of projection values from COVID-19 and non-COVID-19 abstracts onto the first PC, indicating a shared pattern of variance, i.e. both groups appear to discuss similar questions, approaches, and techniques using similar vocabulary within this pattern.

**Figure 1a, 1b (from left to right).**
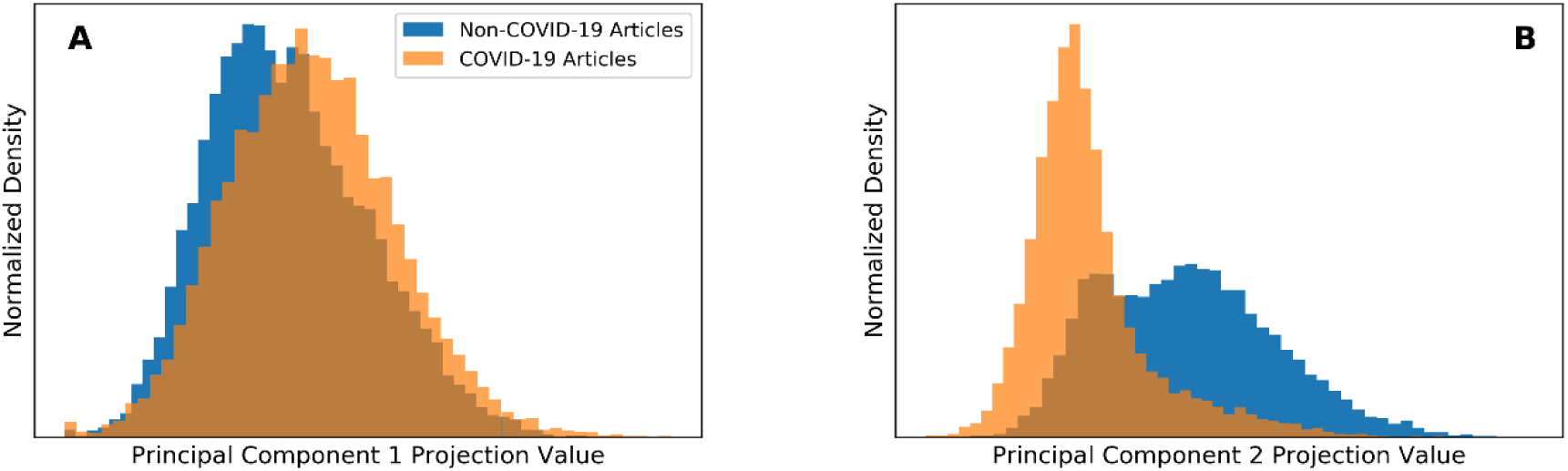
Distribution of COVID-19 (orange) and non-COVID-19 (blue) abstracts along the top two PCs. Panel A shows PC1, and Panel B – PC2. PC1 does not effectively distinguish between COVID-19 and non-COVID-19 abstracts. PC2 shows distinct distributions between COVID-19 and non-COVID-19 abstracts, indicating distinct vocabularies used in these abstracts.

The patterns that successfully differentiated between the two groups were beneath the first PC, within the second PC, where the projection value distributions presented distinguishing patterns (Figure 1b). Our interpretation of this PC relied on identifying terms that had values with the greatest magnitude (Supplemental Information 4; Supplemental Information 5). Ultimately, the figures below indicate that while variance among non-COVID-19 abstracts (blue) stretched over much of the second PC, projection values of COVID-19 abstracts (orange) were concentrated in a smaller area, reflecting the narrower scope of COVID-19 abstracts considering that the virus and associated disease have only been studied since December 2019.

When we split the studies into subsets for the three human coronaviruses that have potential for severe infection, we found that the distributions of SARS-CoV and MERS-CoV abstracts in the PC projection space were unique to each virus (Figure 2). SARS-CoV-2 abstracts appeared to share a space in common with both MERS-CoV and SARS-CoV, likely reflecting some shared terminology and possible ongoing attempts to leverage existing knowledge of the other two viruses to learn about SARS-CoV-2. However, SARS-CoV-2 abstracts are much more concentrated among lower projection values. Notably, MERS-CoV and SARS-CoV abstracts were spread more evenly along the second PC, reflecting greater breadth and variation along these PCs that can be attributed to a broader range of studies focused on these pathogens as compared to SARS-CoV-2. This may be in part due to the much longer time that has been spent studying these viruses.

**Figure 2.**
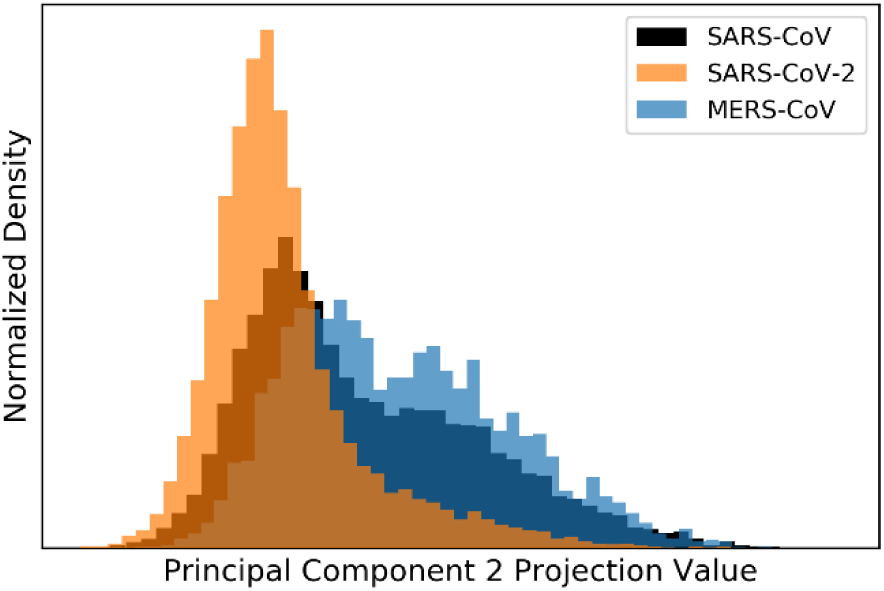
The second PC provides distinct separation of SARS-CoV-2, as well as mild separation between abstracts mentioning the two other human CoVs capable of causing severe illness (SARS-CoV and MERS-CoV).

To identify terms associated with differences between COVID-19 and non-COVID-19 abstracts on PC2, we examined patterns of lemmatized terms from the respective abstracts (Figure 3). The projection values of COVID-19 abstracts on PC2 were lower and associated with emergent COVID-19 clinical-, modeling- or field-based (CMF) research – such as observational, clinical, and epidemiological studies – exemplified by stem terms “patient”, “pandem”, “estim”, and “case”. Words in the opposite direction on PC2 – such as “protein”, “cell”, “bind”, and “express” – can be associated with viral biology and basic disease processes studied in biomolecular laboratories. COVID-19 abstracts were thus mostly associated with research conducted outside of laboratories, e.g., in hospitals, likely reflecting the pandemic reality of data collection alongside (and often secondary to) clinical care.

**Figure 3.**
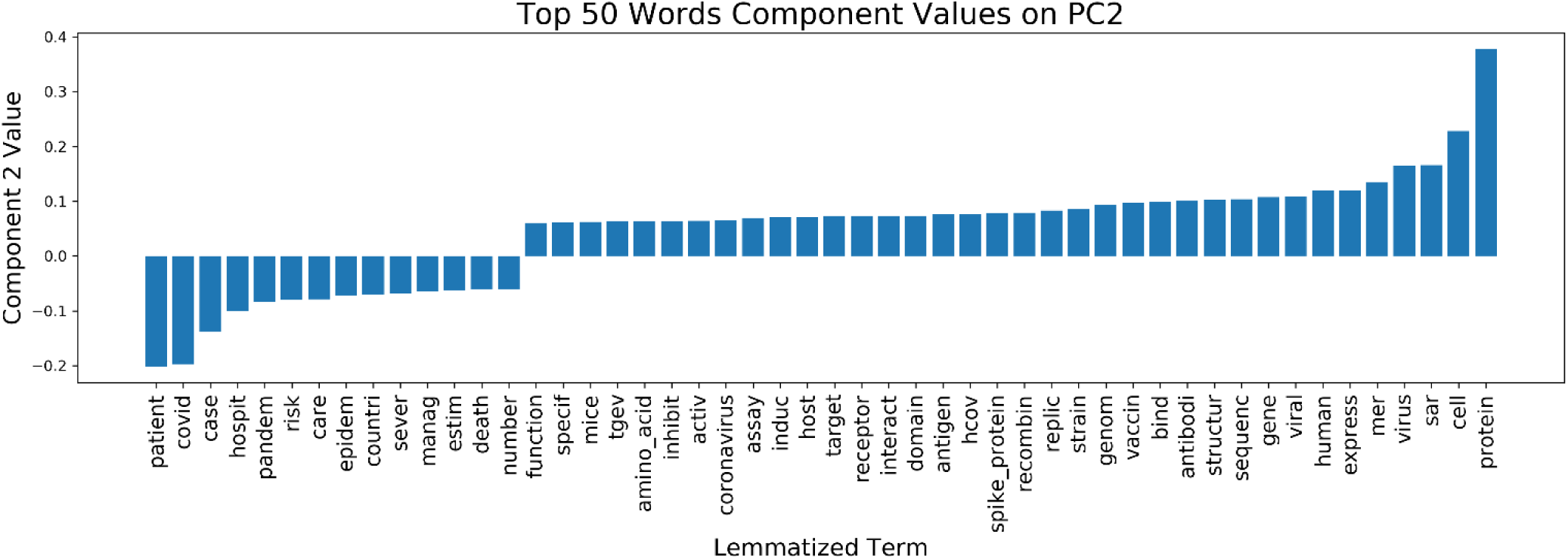
This bar chart displays the values of key lemmatized words on the second PC. Those with greatest magnitude were used for interpretation of the text corpus.

The high-level abstraction reflected by PC2 informed our designation of the extent that COVID-19 research included studies with any CMF design – ranging from epidemiological studies to retrospective reviews of clinical outcomes, case studies, and randomized clinical trials – or laboratory-driven research – including observational microscopy, experimentation with antiviral compounds, derivation of protein structures, and studies of animal or cell culture models. Overall, COVID-19 abstracts appeared more likely to have terms associated with CMF research rather than laboratory studies based on comparisons of distributions for key terms in the COVID-19 and non-COVID-19 abstracts (Figure 4; Supplemental Information 4). This partition along research design for non-COVID-19 and COVID-19 abstracts was also evident in the abstract texts: 90% of the abstracts in the bottom 1% of projection values along the second PC were related to COVID-19; conversely, only 1% of the abstracts in the top 1% were related to COVID-19.

**Figure 4.**
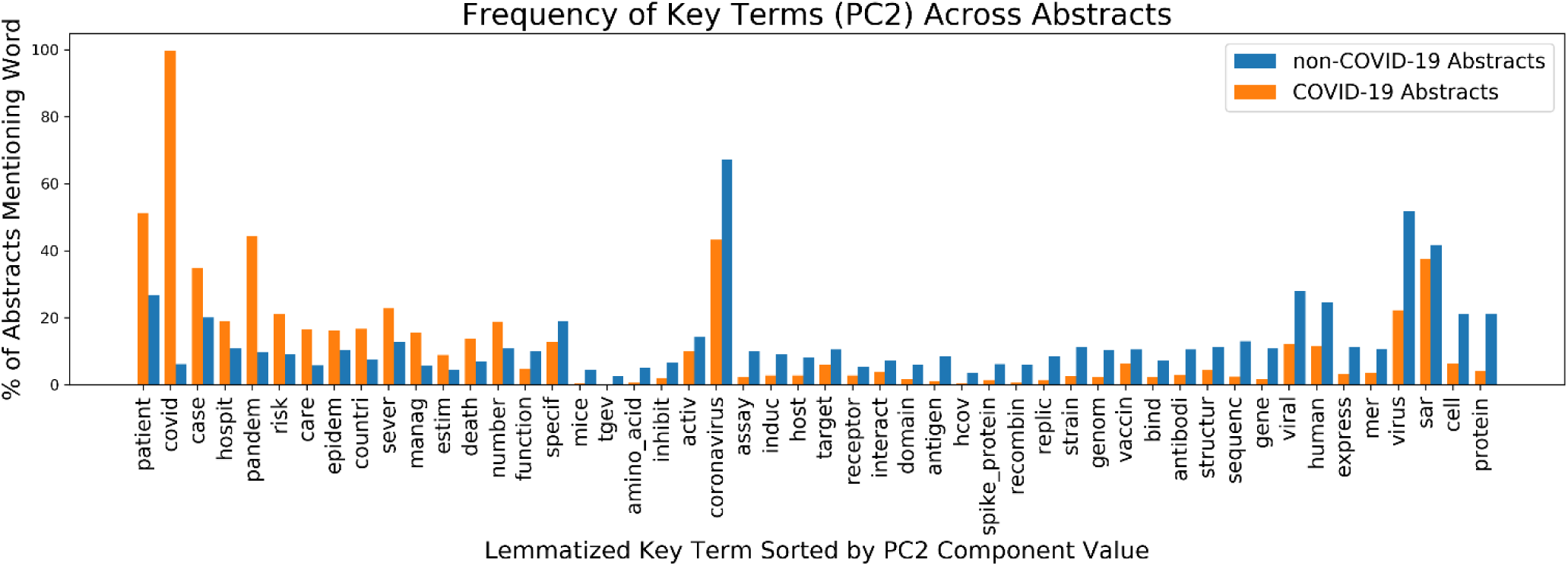
Top 50 key terms are unevenly distributed among the COVID −19 and non-COVID-19 abstracts.

### Topic Modeling Suggests Additional Differences in Specific Research Subareas

Topic modeling helped characterize differences between research topics discussed in COVID-19 and non-COVID-19 abstracts. Results from the LDA model suggested that, similar to the pattern observed in Figure 4, there was clear differentiation between COVID-19 and non-COVID-19 abstracts across 30 topics (Figure 5; Supplemental Information 6). There were five topics in particular – (1) Topic 14: outbreaks’ impact on healthcare services, (2) Topic 15: testing for coronaviruses, (3) Topic 17: epidemic cases and modeling (4) Topic 21: clinical care and therapeutics, and (5) Topic 25: lessons learned for epidemic preparedness – that accounted for 58% of all COVID-19 abstracts and for just 17% of non-COVID-19 abstracts. COVID-19 abstracts were thus disproportionately concentrated in these five topics relative to non-COVID-19 abstracts.

**Figure 5.**
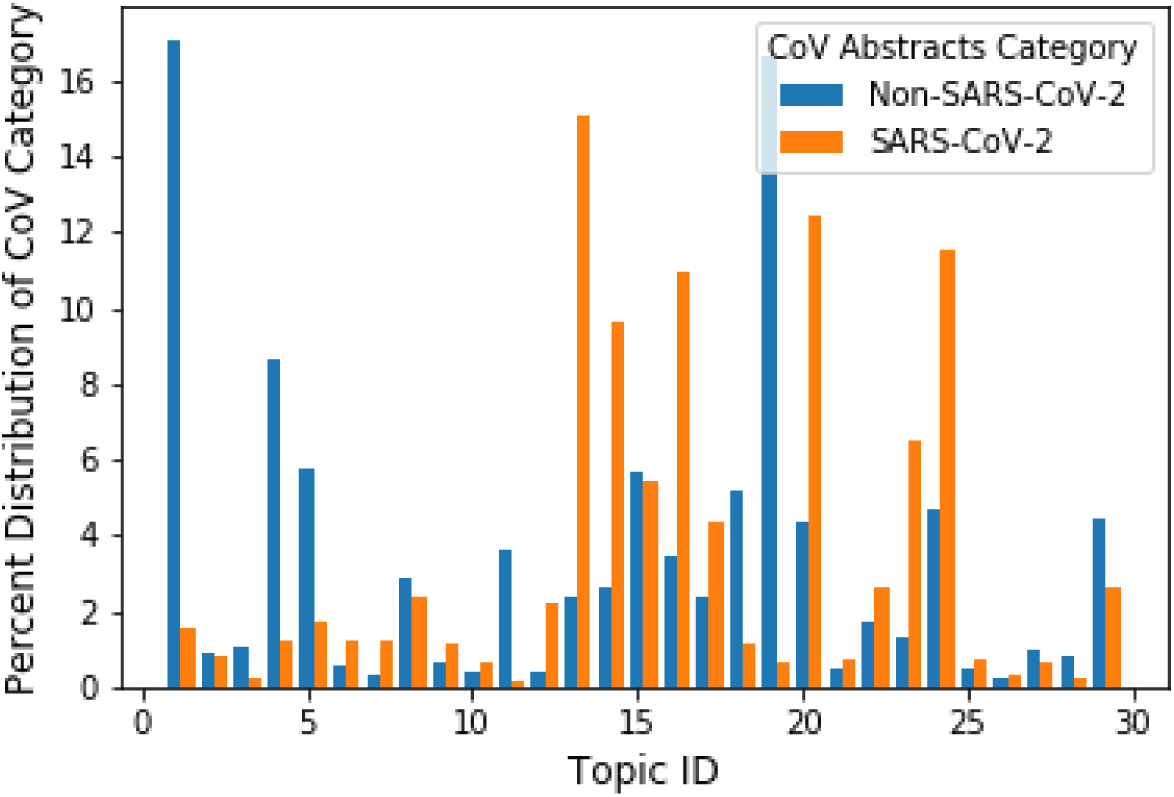
COVID-19 literature is distributed unevenly across the 30 topics.

Across the 30 topics, we grouped several topics into topic families based on internal commonalities (Supplemental Information 6), including (1) updates on the spread of and events related to coronavirus outbreaks (including two subfamilies: general updates vs. public health responses); (2) testing for coronaviruses; (3) clinical care, therapeutics, and the need for vaccinations; and (4) basic microbiological research (which included two subfamilies: a general catch-all subfamily vs. a subfamily specific to pathogenesis and transmission). When divided by topic family, the disparity between COVID-19 and non-COVID-19 research in the first and fourth topic families showed that COVID-19 abstracts appeared to be heavily concentrated on topics that typically included field-based data (the first topic family, on outbreak reporting) and excluded laboratory-based studies (the fourth topic family, on basic microbiology). However, one exception was that COVID-19 was overrepresented in studies on testing (especially diagnostics; the second topic family), which included both the laboratory development of the tests and their field application. (Supplemental Information 7)

### Documents Analyzed Through Machine Learning Highlight Trends Over Time

We also examined the rates of publication and preprint submission for COVID-19 abstracts along PC2 (Figure 6a) and the previously mentioned topic families (Figure 6b). From the beginning of 2020, COVID-19 abstracts tended to have lower projection values for the second PC, reflecting the relatively higher number of CMF studies emerging during the early stages of the pandemic compared to laboratory-based studies.

**Figures 6a and 6b.**
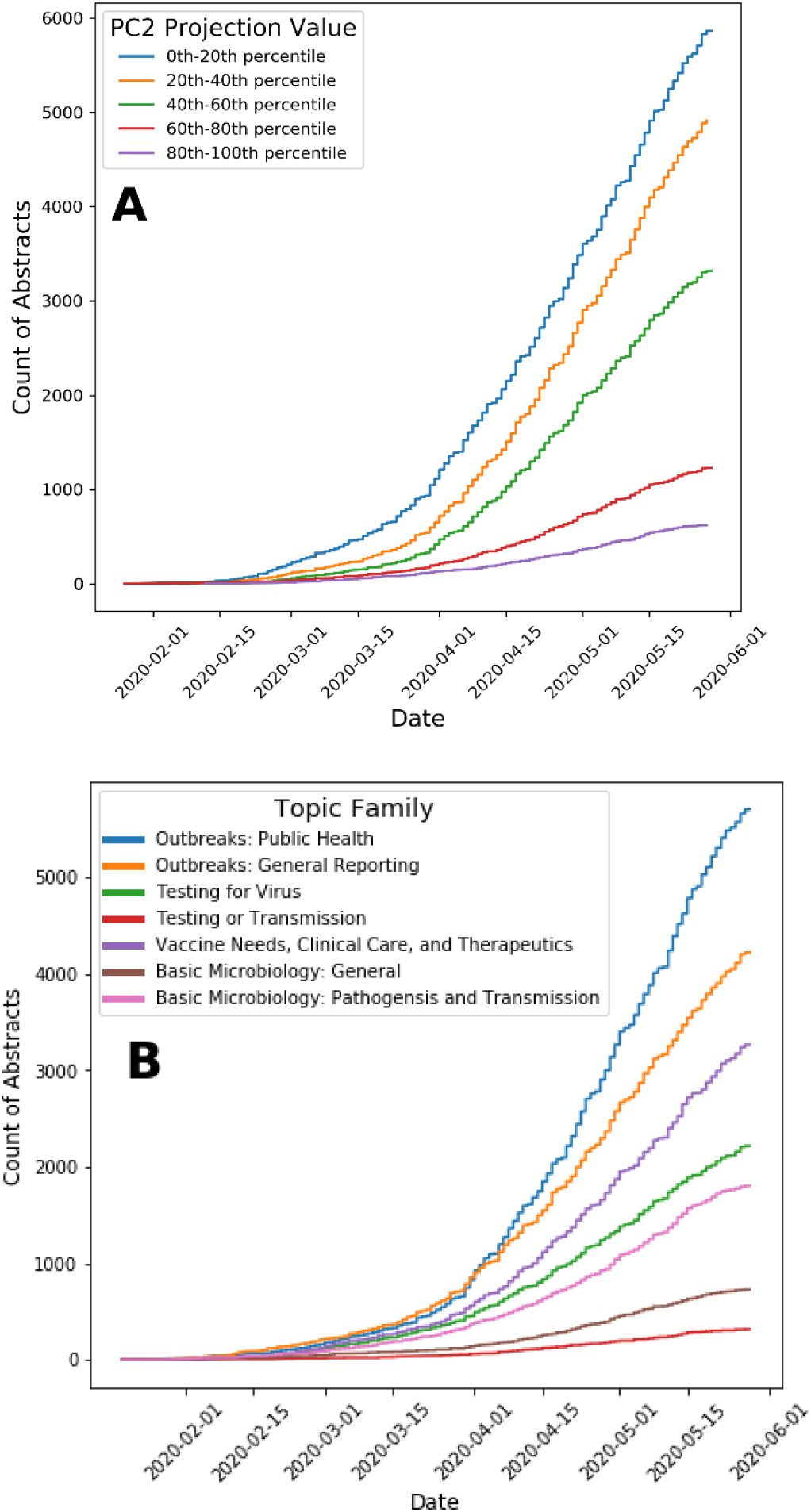
Panel A shows the distribution over time of COVID-19 abstracts with different projection values on the second PC (i.e, those likely reflecting CMF research versus laboratory research) and the different timelines for publication between these groups. Panel B shows COVID-19 research is predominantly focused on outbreak reporting and public health issues.

Likewise, the growth of studies in the different topic families for COVID-19 was unevenly distributed (Figure 6b). From January 2020 through the end of May 2020, publications related to COVID-19 were dominated by studies involving (1) outbreak and responses and (2) patients and healthcare services, similar to the observed faster pace of CMF research in the PCA results. Publications regarding viral mechanisms and biomolecular processes related to SARS-CoV-2 grew at a slower pace.

## Discussion

Our findings demonstrate the utility of our novel NLP-driven approach for determining potential areas of underrepresentation in current research efforts for COVID-19. By applying unsupervised ML methods to CORD-19, we identified overarching key research topics in existing coronavirus and COVID-19-specific abstracts, as well as the distribution of abstracts among topics and over time. Our results support a prior bibliometric study that also found more frequent appearances of epidemiological keywords in COVID-19 research compared to research on other coronaviruses (Hossain, 2020). However, our study presents the unique finding that laboratory-based COVID-19 studies, including those on genetic and biomolecular topics, are underrepresented relative to studies of epidemiological and clinical issues, particularly when compared with the distribution of previous research on other coronaviruses. Furthermore, we developed a framework that improves upon existing studies in two key ways: (1) our method maps connections between abstracts or publications by relying directly on the abstracts’ text in comparison to other bibliometric analyses, including those in other fields, that rely on the analysis of metadata (de Oliveira et al., 2019; Campbell et al., 2010); and (2) our method offers an unsupervised ML-driven approach to splice the data in multiple ways, including adeptly measuring the scope of existing literature, its topical changes over time, and differences from literature on previous pandemics.

The distribution of COVID-19 and non-COVID-19 abstracts from our PCA results suggest that, at the time of writing (CORD-19 dataset release on May 28, 2020), the breadth of published research for COVID-19 is relatively narrow compared to that of published non-COVID-19 studies (Figures 1 and 2). As shown in our results, keywords associated with biomolecular processes (e.g., viral structure, pathogenesis, and host cell interactions) appeared more frequently in non-COVID-19 abstracts than in COVID-19 abstracts. This finding reflects the emergent nature of SARS-CoV-2 and the research community’s struggle to understand it at the molecular level to the same extent as other coronaviruses. Nonetheless, the availability of laboratory studies for other coronaviruses represents an opportunity for generating hypothesis-driven research questions grounded in empirical research.

It is worth noting that researchers may be working under the assumption that biological processes of SARS-CoV-2, including life cycle and interactions with the human host, are comparable to those of SARS-CoV due to their genetic similarity and relatedness (CSG, 2020; Petrosillo et al., 2020; Zhang and Holmes 2020). For example, several prior SARS-CoV studies on host cell entry helped identify the angiotensin converting enzyme 2 (ACE2) protein as a mediator for SARS-CoV-2 infection (Hoffman et al., 2020). Likewise, CD147 and GRP78 proteins have been hypothesized to play a role in cell entry for SARS-CoV-2 based on earlier SARS-CoV and MERS-CoV findings, although additional studies are needed (Wang et al., 2020b; Chen et al., 2005; Ibrahim et al., 2020; Chu et al., 2018). While building upon assumed similarities is an important first step, as work progresses, it becomes increasingly important to identify features that are unique to each virus. However, the scope of literature for biological processes unique to SARS-CoV-2 is currently quite limited, and perhaps even more limited than what our PCA results suggest if most SARS-CoV-2 literature relies heavily on other coronavirus research.

This underrepresentation of studies on biomolecular processes could also be attributed to the rapid worldwide spread of SARS-CoV-2 that occurred within mere months of its emergence, necessitating an unprecedented response from healthcare and public health infrastructures globally. Our PCA results reflect an overwhelming concern regarding the exponential spread of the virus and risks for transmission involved with more frequent appearances of stem terms such as “pandem”, “outbreak”, “estim”, “countri”, “number”, and “risk” in COVID-19 abstracts. This was also supported by our topic modelling results, which indicated that 58% of COVID-19 abstracts fell into just five of 30 topics, generally related to healthcare services, the pandemic’s public health issues, and testing for coronaviruses (Figure 6a, 6b). The more rapid growth of CMF research, relative to laboratory-driven research, mirrors the current response to the pandemic in the United States, where the initial focus on pressing epidemiological and clinical concerns is now followed by interest in experimental investigations, including those of structural mechanisms for host cell entry and possible therapeutic targets.

Overall, our findings reflect a clear divide between COVID-19 and non-COVID-19 abstracts based upon research design; unlike CMF research, laboratory-driven SARS-CoV-2 research is either still underway or has only just been initiated. This can be attributed in part to the fact that laboratory research is often a labor-intensive process within a federally-regulated infrastructure that depends on the availability of timely, project-based funding as well as longer-term funding. Our findings also suggest that the pace of research on SARS-CoV-2 biomolecular processes is potentially insufficient given the global threat posed by the virus (Figure 6a, 6b). This lag may adversely impact the development of antivirals and other therapeutic interventions, adding strain to already overwhelmed healthcare systems. Furthermore, these trends raise questions about the readiness of institutions supporting the research community in times of extraordinary stress. Previous experiences with global pandemics, such as H1N1, have resulted in various policy recommendations (French et al., 2009) to maintain and enhance readiness in laboratory-based research, and analysis on the effectiveness of recommendations arising from these experiences may be worthwhile.

While PCA identified a prominent pattern that differentiated between COVID-19 and non-COVID-19 literature, the topic families derived from LDA refined our understanding of knowledge gaps and research needs in COVID-19 literature by delineating specific research areas. This included an underrepresentation of studies on basic microbiological examination of SARS-CoV-2, including its pathogenesis and transmission. Research on these issues is published at a slower pace than CMF studies (e.g., those on clinical topics, outbreak response, and statistical reporting) and research on testing (Figure 6b). Even when compared with the distribution of non-COVID-19 research, COVID-19 research was more heavily focused on topics within the CMF realm (Supplemental Information 7).

We recognize that the number of abstracts in each of these topics does not necessarily represent scientific progress made in these areas, but they *do* reflect the pace of research and potential availability of public knowledge. This indicates either a mismatch between the level of effort in these issues and the urgency of work or time lags inherent to these fields that constrain the responsiveness of the scientific community. Increased and consistent funding of emerging pathogens research, including support of basic research even when there is no immediate threat of an outbreak, would allow us to maintain a proactive posture in accumulating available knowledge rather than over-reliance on reactivity.

These conclusions must be caveated by several limitations that must be acknowledged. First, while CORD-19 includes a vast quantity of coronavirus-related publications, it potentially omits relevant literature from other databases, such as the Social Science Research Network (SSRN) or arXiv (a preprint server for studies in mathematics, computer science, and quantitative biology, among other topics). This may have constrained the representativeness of our analysis on COVID-19 literature, thus affecting the external validity of our findings. Second, analyzing abstracts inherently excludes ongoing research efforts because not all relevant studies are publicly available or have released preprints. Third, the number of publications does not directly represent progress in research areas. Fourth, the high-level trends we observed through our unsupervised ML approaches may not completely align with how researchers identify and process specific research topics. The counts of words in DTMs informing the ML algorithms may not directly capture the ideas researchers are trying to convey and may therefore gloss over nuances in the literature.

Yet these four limitations are somewhat mitigated by both the nature of the data sources and the needs of the research community. For the first, the excluded sources (SSRN and arXiv) heavily focus on research within the CMF arena, indicating that if anything, our conclusions on the rapid pace of CMF COVID-19 research (versus lab-based research) are conservative. For the second, existing research pipelines have been accelerated in the pandemic, especially with the proliferation of pre-print services. This reduces the lag between the discovery of knowledge and the availability of an abstract to ingest in our data pipeline. Third, the number of publications in each area may imply a relative difference in research productivity for different topics, and thus may still serve as a proxy for indicating such progress or the attention given to specific issues.

And finally, our ML-based method offers the chance to quickly review large quantities of text at scale and highlight underlying trends. Both this speed and this scale are crucial to informing time-sensitive decisions on policy and priorities to facilitate the most impactful research.

Complex public health problems like the ongoing COVID-19 pandemic require researchers to maintain robust knowledge on pathogenic threats, including efforts dedicated to emerging or currently neglected pathogens. Our ML-based study offers insights into potential areas for future research opportunities and funding investments that build upon what is already known about coronaviruses. We offer a conceptual framework that can be applied to other emergent or neglected pathogens that have the potential to become a pandemic threat, such as highly pathogenic avian influenza H5N1 (Kilpatrick et al., 2006; Kissler et al., 2019), enabling researchers to maintain a proactive preparedness posture.

## Availability of Data and Materials

The data of CORD-19 is available to download from <u>here</u>. All the code is free for download from <u>GitHub here</u>.

## Acknowledgements

This project is part of the COVID-19 Dispersed Volunteer Research Network (COVID-19-DVRN) led by Dr. Maimuna Majumder and Dr. Angel Desai. Dr. Majumder is supported by a grant from the National Institutes for Health (NIH) (T32HD040128) and is a faculty member at both Harvard Medical School and Boston Children’s Hospital’s Computational Health Informatics Program (CHIP).

Dr. Helen Piontkivska is also supported by a grant from the NIH (R21AG064479-01). The funders for this grant had no role in the design of the study and collection, analysis, and interpretation of data and in writing the manuscript.

The authors thank Shagun Gupta for her thoughtful feedback on an earlier version of the manuscript.

## Author Contributions

Conceptualization: Anhvinh Doanvo (A.D.), Xiaolu Qian (X.Q.), Divya Ramjee (D.R.), Helen Piontkivska (H.P.), Angel Desai (A.D.2), Maimuna Majumder (M.M.); Data Curation, Methodology, Software, and Visualization: A.D. and X.Q.; Formal Analysis and Investigation: A.D., X.Q., D.R., and H.P.; Writing – Original Draft: A.D., D.R., X.Q., and H.P.; Writing – Review & Editing: A.D., D.R., H.P., X.Q., A.D.2, M.M.

## Footnotes

1 https://github.com/COVID19-DVRN/8-AI-Mapping-of-Relevant-Coronavirus-Literature

