## Supplemental Information 1 for "Machine Learning Maps Research Needs in COVID-19 Literature"

***Supplemental Information 1. Word Counts in Document-Term Matrices Serve as the Text's Computable Features***

The smallest meaningful unit of semantic information in human language is a word. Therefore, we can infer that documents with very different words—perhaps shown through different frequencies of specific terms—discuss different topics; documents with similar frequencies for the same terms are likely focused on similar topics. This quantitative information feeds easily into classical machine learning algorithms, and so we captured it through document-term matrices (DTMs). Each cell in a DTM is filled by a metric for the frequency of a term (a column) in a specific document (a row). The DTM that we fed into PCA was a term frequency-inverse document frequency matrix, which down-weights terms that are more common across all products and thus aren't likely to be good differentiators between abstracts. LDA, on the other hand, expects simple term counts.

Since the words represent the feature space of DTMs, we took care to identify terms in a meaningful manner. Prior to computing the DTMs, we removed all punctuation and numbers from the text and lemmatized the remaining words so that words with the same stem are consolidated. This reduced the noise in the dataset and enhanced the consistency between machine-derived metrics and their semantic meaning. We also leveraged existing natural-language-processing packages (Gensim) to identify potentially useful word pairs, or "bigrams", as terms to feed into the DTMs.
