## Supplemental Information 2 for "Machine Learning Maps Research Needs in COVID-19 Literature"

*Supplemental Information 2. COVID-19-related Keywords for Filtering Subsetted Abstracts*

| Item | Value(s) | Description and rationale |
|---|---|---|
| **General search terms** | Case sensitive: MERS<br>Not case sensitive: "covid-19", "coronavirus", "corona virus", "2019-ncov", "sars-cov", "mers-cov", "severe acute respiratory syndrome", "middle east respiratory syndrome" | • The presence of these search terms in an abstract indicated that the abstract was relevant to the study<br>• Mentioning these terms in an abstract made it more likely that a coronavirus was central to the research |
| **Search terms for COVID-19** | "COVID-19", "COVID", "2019-nCoV", "SARS-CoV-2" (case sensitive) | • The presence of these terms in an abstract indicated that it was relevant to COVID-19 |
| **Search terms for MERS** | Case sensitive: MERS<br>Not case sensitive: "middle east respiratory" | • The presence of these terms in an abstract indicated that it was relevant to MERS-CoV |
| **Search terms for SARS** | Case sensitive: SARS<br>Not case sensitive: "severe acute respiratory syndrome" | • The presence of these terms in an abstract indicated that it was relevant to SARS-CoV |
| **Number of topics** | 30; identified through manual review of multiple topic models. | • Neither perplexity nor coherence metrics yielded optimal configurations without an excessive number of topics.<br>• Manual review for relative mutual exclusivity and tightness of topics set the criteria |
