## Supplemental Information 3 for "Machine Learning Maps Research Needs in COVID-19 Literature"

***Supplemental Information 3. Randomized PCA Dramatically Speeds Computation***

The complexity of principal components analysis through singular value decomposition (SVD) scales with an order of $O(max(m,n) * min(m,n)^2)$, or $O(m * n^2)$ when $m$ is large, and where $m$ and $n$ are the number of observations (documents) and features (unique words) respectively. With nearly 10,000 articles mentioning coronavirus-related terms in their abstracts and tens of thousands of unique words, SVD computations can take some time. We accelerated this step by using randomized SVD, which has an order of complexity of just $O(mn\ log(k))$, where $k$ is the number of principal components computed. Indeed, this enabled our SVD calculations to proceed almost instantaneously. And while there is some randomness associated with results from randomized SVD, existing literature indicates that its output converges super-exponentially to the true output of SVD with additional iterations. (Halko, Martinsson and Tropp, 2011)
