## Supplemental Information 4 for "Machine Learning Maps Research Needs in COVID-19 Literature"

***Supplemental Information 4. Distribution of the Top 50 Key Terms Separating COVID-19 Abstracts from non-COVID-19 Abstracts along Principal Component 2 (shown in Figure 3)***

| **Lemmatized Word** | **Component Value** | **Percentage of COVID-19 Abstracts** | **Percentage of non-COVID-19 Abstracts** |
|---|---|---|---|
| patient | -0.20184 | 51.3 | 26.7 |
| covid | -0.19725 | 99.8 | 6.1 |
| case | -0.13793 | 34.9 | 20.1 |
| hospit | -0.09987 | 18.9 | 10.8 |
| pandem | -0.08412 | 44.4 | 9.7 |
| risk | -0.07939 | 21.2 | 9.1 |
| care | -0.0786 | 16.6 | 5.8 |
| epidem | -0.07234 | 16.1 | 10.4 |
| countri | -0.07022 | 16.7 | 7.6 |
| sever | -0.06779 | 22.9 | 12.8 |
| manag | -0.06461 | 15.5 | 5.6 |
| estim | -0.06186 | 8.9 | 4.5 |
| death | -0.06158 | 13.8 | 7 |
| number | -0.06069 | 18.8 | 10.8 |

| | | | |
|---|---|---|---|
| function | 0.060604 | 4.8 | 10.1 |
| specif | 0.06181 | 12.8 | 18.9 |
| mice | 0.062183 | 0.4 | 4.5 |
| tgev | 0.06324 | 0 | 2.6 |
| amino_acid | 0.063691 | 0.6 | 5.1 |
| inhibit | 0.063901 | 2 | 6.6 |
| activ | 0.064333 | 10.1 | 14.2 |
| coronavirus | 0.065021 | 43.3 | 67.1 |
| assay | 0.069902 | 2.3 | 10 |
| induc | 0.070871 | 2.7 | 9.1 |
| host | 0.071237 | 2.7 | 8.2 |
| target | 0.072913 | 5.9 | 10.5 |
| receptor | 0.07325 | 2.7 | 5.4 |
| interact | 0.07327 | 3.9 | 7.3 |
| domain | 0.073451 | 1.6 | 6 |
| antigen | 0.075788 | 1 | 8.4 |
| hcov | 0.076117 | 0.4 | 3.5 |

| | | | |
|---|---|---|---|
| spike_protein | 0.078466 | 1.5 | 6.1 |
| recombin | 0.079413 | 0.6 | 6 |
| replic | 0.082887 | 1.5 | 8.5 |
| strain | 0.085891 | 2.5 | 11.3 |
| genom | 0.094031 | 2.2 | 10.3 |
| vaccin | 0.097755 | 6.2 | 10.5 |
| bind | 0.099378 | 2.3 | 7.3 |
| antibodi | 0.100884 | 2.9 | 10.5 |
| structur | 0.103286 | 4.3 | 11.2 |
| sequenc | 0.104208 | 2.4 | 13 |
| gene | 0.108125 | 1.6 | 10.8 |
| viral | 0.10938 | 12.2 | 27.9 |
| human | 0.119375 | 11.5 | 24.6 |
| express | 0.119808 | 3.2 | 11.3 |
| mer | 0.134674 | 3.6 | 10.6 |
| virus | 0.164846 | 22.2 | 51.8 |
| sar | 0.165963 | 37.4 | 41.7 |

| | | | |
|---|---|---|---|
| cell | 0.228506 | 6.3 | 21 |
| protein | 0.378095 | 4.1 | 21.3 |
