## Supplemental Information 5 for "Machine Learning Maps Research Needs in COVID-19 Literature"

***Supplemental Information 5. Examples of Abstracts Identified as Either COVID-19 or non-COVID related***

| **Relationship with COVID-19** | **Abstract Title** |
|---|---|
| Related to COVID-19<br><br>(Bottom 1% of SecondPrincipal Component) | Recommendations for standardized management of CML patients in the core epidemic area of COVID-19 (Wang et al., 2020c) |
| | Transmission risk of patients with COVID-19 meeting discharge criteria should be interpreted with caution (Su, 2020) |
| | COVID-19 in a Designated Infectious Diseases Hospital Outside Hubei Province, China (Cai, 2020) |
| Unrelated to COVID-19<br><br>(Top 1% of Second Principal Component) | Characterization of the expression and immunogenicity of the ns4b protein of human coronavirus 229E. (Chagnon, 1998) |
| | Severe acute respiratory syndrome coronavirus nucleocapsid protein expressed by an adenovirus vector is phosphorylated and immunogenic in mice. (Zakhartchouk, 2005) |
| | Molecular cloning and expression of a spike protein of neurovirulent murine coronavirus JHMV variant cl-2. (Taguchi, 1992) |
