## Supplemental Information 6 for "Machine Learning Maps Research Needs in COVID-19 Literature"

*Supplemental Information 6. Topic Families Across COVID-19 and non-COVID-19 Abstracts*

| ID | Topic Title | Topic Family | COVID-19 Count (percent) | Total (percent) |
|---|---|---|---|---|
| 1 | Treatment and patient care for COVID-19 | Vaccine needs, patient care, and treatments | 250 (1.4%) | 338 (1%) |
| 2 | Biomolecular study of coronaviruses | Microbiology (general) | 287 (1.6%) | 3165 (9%) |
| 3 | Infection by coronavirus | Microbiology (transmission) | 149 (0.8%) | 311 (0.9%) |
| 4 | Porcine coronavirus microbiology and infection | Microbiology (general) | 37 (0.2%) | 199 (0.6%) |
| 5 | Infection by coronavirus | Microbiology (transmission) | 226 (1.2%) | 1695 (4.8%) |
| 6 | Public health issues of SARS transmission | Outbreaks (public health) | 305 (1.7%) | 1274 (3.6%) |
| 7 | Outbreaks in different countries | Outbreaks (general coverage) | 231 (1.3%) | 336 (1%) |
| 8 | Death and mortality due to COVID-19 | Outbreaks (general coverage) | 226 (1.2%) | 284 (0.8%) |
| 9 | Human infection by coronaviruses | Microbiology (transmission) | 430 (2.3%) | 911 (2.6%) |
| 10 | Testing and infection of COVID-19 | Testing (mixed with transmission) | 210 (1.1%) | 321 (0.9%) |
| 11 | Impact of COVID-19 on community services | Outbreaks (general coverage) | 125 (0.7%) | 208 (0.6%) |
| 12 | Infection by MERS-CoV | Microbiology (transmission) | 36 (0.2%) | 660 (1.9%) |

| | | | | |
|---|---|---|---|---|
| 13 | General COVID-19 pandemic coverage | Outbreaks (general coverage) | 432 (2.3%) | 498 (1.4%) |
| 14 | COVID-19's impact on healthcare services | Outbreaks (public health) | 2666 (14.5%) | 3050 (8.6%) |
| 15 | Clinical testing and COVID-19 symptoms | Testing | 1750 (9.5%) | 2201 (6.2%) |
| 16 | Transmission and infection among coronaviruses | Microbiology (transmission) | 980 (5.3%) | 1903 (5.4%) |
| 17 | Modeling, statistics, and investigation of epidemic | Outbreaks (general coverage) | 2006 (10.9%) | 2580 (7.3%) |
| 18 | Drug studies and need for vaccines | Vaccine needs, patient care, and treatments | 837 (4.5%) | 1214 (3.4%) |
| 19 | Study of coronavirus genomes | Microbiology (general) | 201 (1.1%) | 1054 (3%) |
| 20 | Biomolecular study of coronaviruses | Microbiology (general) | 123 (0.7%) | 2931 (8.3%) |
| 21 | Treatment and patient care for coronaviruses | Vaccine needs, patient care, and treatments | 2234 (12.1%) | 2958 (8.4%) |
| 22 | Children infected by COVID-19 | Outbreaks (general coverage) | 125 (0.7%) | 216 (0.6%) |
| 23 | Clinical testing for COVID-19 | Testing | 475 (2.6%) | 767 (2.2%) |
| 24 | COVID-19 cases and deaths reported | Outbreaks (general coverage) | 1190 (6.5%) | 1408 (4%) |
| 25 | Lessons learned for epidemic preparedness | Outbreaks (public health) | 2038 (11.1%) | 2801 (7.9%) |

| | | | | |
|---|---|---|---|---|
| 26 | Outbreak and public health response to COVID-19 | Outbreaks (public health) | 138 (0.7%) | 221 (0.6%) |
| 27 | Biomolecular study of coronaviruses | Microbiology (general) | 56 (0.3%) | 89 (0.3%) |
| 28 | Testing and transmission of COVID-19 | Testing (mixed with transmission) | 121 (0.7%) | 296 (0.8%) |
| 29 | Biomolecular study of coronavirus strains | Microbiology (general) | 51 (0.3%) | 191 (0.5%) |
| 30 | Outbreaks and public health responses | Outbreaks (public health) | 477 (2.6%) | 1200 (3.4%) |
