## Supplemental Information 7 for "Machine Learning Maps Research Needs in COVID-19 Literature"

***Supplemental Information 7. The percentage of all COVID-19 abstracts in each of the broad research topic.***

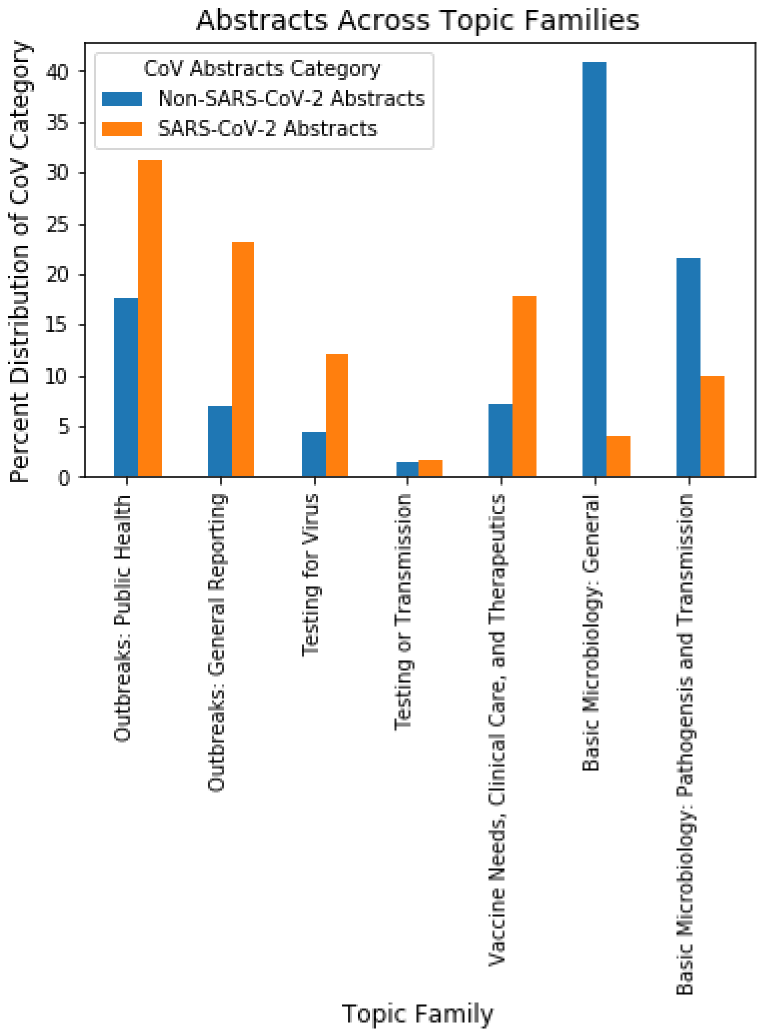
